# Exogenous attention facilitates performance through dissociable sensitivity and bias mechanisms

**DOI:** 10.1101/380659

**Authors:** Vishak Sagar, Ranit Sengupta, Devarajan Sridharan

## Abstract

Selective attention enables prioritizing the most important information for differential sensory processing and decision-making. Here, we address an active debate regarding whether attention reflects a unitary phenomenon or involves the operation of dissociable sensory enhancement (perceptual sensitivity) and decisional gating (choice bias) processes. We developed a multialternative task in which participants detected and localized orientation changes in gratings at one of four spatial locations. Exogenous attention cues (high contrast flashes) preceded or followed the change events in close temporal proximity. Analysis of participants’ behavior with a multidimensional signal detection model revealed markedly distinct effects of exogenous cueing on perceptual sensitivity and choice bias. Whereas sensitivity enhancement was localized to the stimulus proximal to the exogenous cue, bias enhancement occurred even for distal stimuli in the cued hemifield. Modulations of sensitivity and bias were uncorrelated at both cued and uncued locations. Finally, exogenous cueing produced reaction time benefits only at the cued location and costs only at locations contralateral to the cue. These disparate effects of exogenous cueing on sensitivity, bias and reaction times could be parsimoniously explained within the framework of a diffusion-decision model, in which the drift rate was determined by a linear combination of sensitivity and bias at each location. Exogenous cueing effects on sensitivity and bias differed systematically from previously reported effects of endogenous cueing. We propose that the search for shared neural substrates of exogenous and endogenous attention would benefit from investigating neural correlates of their component sensory and decisional mechanisms.

**Significance statement:** When we voluntarily direct attention “endogenously”, we are able to better perceive stimuli at the attended location (sensitivity), and to prioritize information from that location for guiding behavioral decisions (bias). But when a salient stimulus, such as a flash of lightning, captures our attention “exogenously”, does it also produce these same effects? To answer this question, we designed a multiple alternative task in which task events occurred in close conjunction with salient exogenous cues (high contrast flashes). We discovered that exogenous attention enhanced both sensitivity and bias for cued stimuli, but each of these changes followed distinct spatial patterns across locations. Our results provide novel insights into component processes of exogenous attention and motivate the search for their neural correlates.

## Introduction

Attention is an essential cognitive capacity for adaptive survival (1, 2). Given the high energy costs (3) associated with neural information processing, the capacity for selective attention permits efficient utilization of the brain’s neural resources by enabling selective processing of task-relevant information while filtering out irrelevant information (4, 5). Selection of information from relevant locations can occur either by endogenous engagement of attention, based on task-relevant goals (e.g. monitoring a traffic light), or by exogenous capture of attention by salient sensory events (e.g. a flash of lightning). In either case, once engaged, attention enables efficient selection through at least one of two mechanisms: i) by enhancing the processing of attended sensory information, often at the cost of unattended information, and ii) by prioritized gating of attended information for guiding perceptual decisions (6–13). These distinct effects of attention can be separately quantified with signal detection theory (SDT), a highly successful framework for the analysis of behavior (14, 15). In conventional SDT, perceptual decision-making is modeled with a latent variable model comprising two parameters: i) perceptual sensitivity (d’) and ii) choice bias (b) (Fig. 1C). While a change in sensitivity reflects the improvement in signal-to-noise for the attended information, a change in bias reflects the prioritized gating of attended information for decisions (16–18).

**Figure 1.**
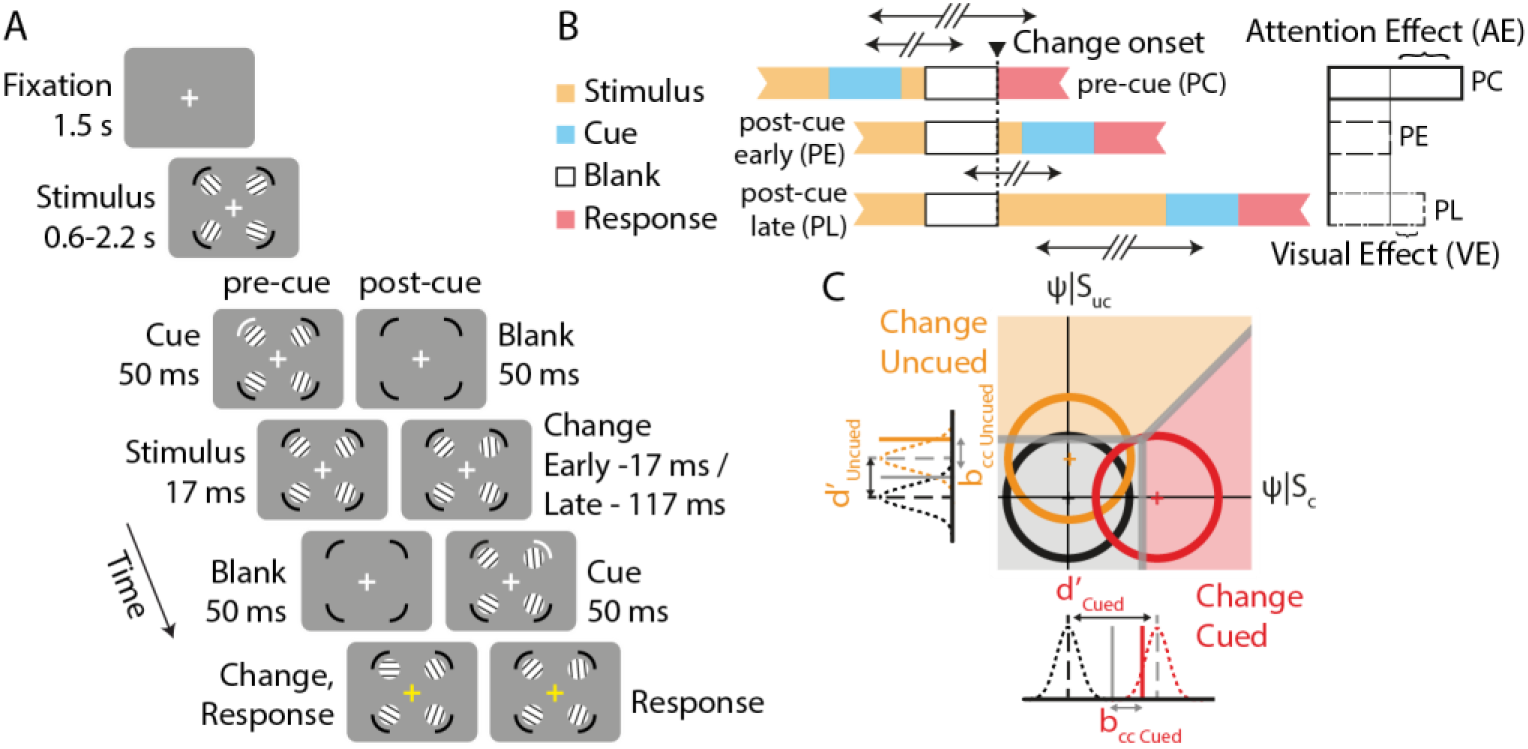
Multialternative change detection task with exogenous cueing. **A**. Change detection task protocol: After an initial fixation (1.5 s), four circular gratings appeared, one in each visual quadrant for a variable duration (exponentially distributed between 0.6-2.2 s). In the pre-cue (PC) trials, an exogenous cue (50 ms flash of cue pedestal) occurred adjacent to any one of the four gratings. After 17 ms the gratings were blanked for 50 ms. When the gratings reappeared, any one or none of the four gratings changed in orientation. Subjects were required to detect and localize the change. In post-cue trials, the exogenous cue was presented at two different latencies (early, PE or late, PL) following grating reappearance (timing described in panel B). In each case, the cue location was independent of change location. PC, PE and PL trials were psuedorandomized and interleaved. **B**. (Left) Schematics showing the timing of exogenous cue (blue) relative to grating reappearance (dashed vertical line) for the three trial types. The post-cue, in PE trials, and the pre-cue, in PC trials, were presented symmetrically about (67 ms after or before) a time point coincident with the centre of the blank epoch, about which information relevant to detecting the change occurred (arrows with two slashes). Similarly, the post-cue, in PL trials, and the pre-cue, in PC trials, were presented symmetrically about (117 ms after or before) a time point 25 ms after the reappearance of the gratings (arrows with three slashes). The exogenous cue is expected to produce both attentional and visual effects in PC trials, visual effects, but not attentional effects, in PE trials, and possibly weak visual effects in PL trials. (Right) Attention effects (AE) of the cue are computed by subtracting behavioral metrics on PE trials from those in PC trials, whereas visual effects (VE) of the cue are computed by subtracting corresponding metrics in PL trials from PE trials. **C**. Schematic of a two-dimensional m-ADC signal detection model. Decision variables at each location are represented along orthogonal decision axes in a multidimensional perceptual space. Circles: Contours of multivariate Gaussian decision variable distributions. Black: noise distribution; red and orange: signal distribution for “change” trials at the cued and uncued locations, respectively. Thick gray lines: Decision surfaces for the multialternative decision. Gray, red and orange shading: Decision zones corresponding to no-change, change at cued location and change at uncued location, respectively. Univariate Gaussians along the axes represent the marginal decision variable distributions at each location.

Here, we study the effect of exogenous visuospatial attention on perceptual sensitivity and decisional bias processes. While there exists an extensive literature on endogenous visuospatial attention (19–21) in a variety of behavioral task contexts, exogenous attention effects have been more challenging to study. This disparity is partly because exogenous attentional processes are transitory, and automatic, operating on fast timescales of a few hundred milliseconds, compared to endogenous attention, which can be usually sustained over several seconds (22, 23). Furthermore, when sensory events that engage exogenous attention are not directly relevant to an ongoing task, sensory processing of the triggering event itself can engage neural resources and interfere with task-related information processing, confounding the approaches for studying the attentional effects in isolation.

Previous studies on exogenous cueing of attention have reported systematic effects on reaction times at the cued location (24–26), as well as behavioral accuracy (27–30). For example, studies that investigated cueing effects on contrast sensitivity by having subjects detect changes in stimulus contrast (31, 32), reported that exogenous cueing consistently facilitated subjects’ ability to detect signals (33) and improved their reaction times (34). Despite these previous reports, little is known about whether these effects of exogenous cueing are mediated through sensitivity or bias changes. Do exogenous cues influence perceptual sensitivity, choice bias, or both? Are these sensitivity and bias effects of exogenous cueing coupled or dissociable? Are the sensitivity and bias effects spatially localized, in proximity to the exogenous cue, or do they impact distal spatial locations also? And, can the sensitivity and bias effects be mechanistically linked to accuracy and reaction time effects of exogenous cueing?

To answer these questions, we designed a multi-alternative change detection task with exogenous spatial cueing (“high contrast” flash, Fig. 1A). The task required subjects to compare sensory evidence across multiple spatial locations to make a single detection and localization decision on each trial. In contrast to previous response probe paradigms (35), our task design permitted measuring spatial bias, across locations, for localizing the change. Because a salient exogenous cue also suppresses visual processing of stimulus information, we designed two post-cue control conditions which permitted isolating pre-cueing attentional effects from visual interference effects of the cue. Subjects’ behavior was analyzed with a recently developed multidimensional signal detection model (36–38) that permitted quantifying exogenous cueing effects on sensitivity and bias at each location. The results reveal key dissociations between sensitivity and bias changes induced by exogenous cueing, and provide mechanistic insights into how exogenous cueing systematically facilitates behavioral accuracies and reaction times by modulating sensitivity and bias.

## Results

### Quantifying attentional and visual effects of exogenous cueing with a multialternative decision task

We investigated the effects of exogenous cueing of attention on perceptual processing and decision-making. Specifically, we investigated how exogenous cueing modulates perceptual sensitivity and choice bias at cued and uncued locations, and whether these effects were coupled or dissociable from one another.

Subjects (n=42) performed a multi-alternative change detection task with exogenous cueing (see Methods for details). Each trial began with a display of 4 stimulus gratings with different orientations, one in each visual quadrant. After an unpredictable delay, an exogenous cue (high contrast arc) appeared briefly (50 ms) adjacent to one of the four stimuli (Fig. 1A), with equal probability across locations (25%). 17 ms following the offset of the cue, the screen was blanked for 50 ms, after which the gratings reappeared. Following reappearance, any one of the four gratings, or none, changed in orientation; the change could occur at any location, independently of the cued location, rendering spatial information pertaining to the cue irrelevant to the task. At the end of each trial, subjects reported one of five possible responses -- orientation change in one of four stimulus gratings, or no change at all -- with one of five distinct key presses (SI Fig. S1A).

To account for sensory interference by the exogenous cue on visual processing and decision-making (“visual interference” effects), we also designed two additional, control conditions involving post-cueing. Figure 1A-B illustrates the experimental protocol for all three trial types: i) the pre-cue (PC) trial in which the exogenous cue occurred 67 ms before the initial grating offset, ii) the “early” post-cue (PE) trial in which the exogenous cue occurred 17 ms after the final grating onset, and iii) the “late” post-cue (PL) trial in which the exogenous cue occurred 117 ms after the final grating onset. The PE trial was designed so that the pre-cue and post-cue were symmetrically distributed about a time point coincident with the centre of the blank epoch; note that information relevant to localizing the change (grating orientation before versus after the change) occurred on either side of the blank epoch (Fig. 1B). Similarly, the PL trial was designed so that the pre-cue and post-cue would be symmetrically distributed about a time point 25 ms after the onset of the potential change (Fig. 1B). Each of these trial types occurred in a pseudorandom sequence in equal proportions (1/3rd each) across trials.

For each trial type (PC, PE, PL), subject’s responses were aggregated into a 5×5 stimulus-response contingency table, with each row and column representing, respectively, change events and responses at locations relative to the cued location (Fig. 2A, SI Fig. S1D). These are, in order: (i) the cued location (Cu), (ii) the location in an adjacent quadrant, ipsilateral (Ip) or (iii) contralateral (Co) to the cue, and (iv) the location in the quadrant diagonally opposite to the cue (Op). The behavioral responses in the 5×5 contingency table were analyzed with a multidimensional signal detection model, the m-ADC model (36–38), to quantify psychophysical parameters (sensitivity and bias) at each location. In Figure 1C we show a schematic of a two-dimensional (2-ADC) model, for illustrative purposes, although these data were analyzed with a four-dimensional (4-ADC) model (see Methods). Occasionally, we report metrics average across uncued locations in the quadrants contralateral to the cue (Co and Op locations, termed Co_av_ metrics) or averaged across all uncued locations (Ip, Co, and Op, termed Uc_av_ metrics).

**Figure 2.**
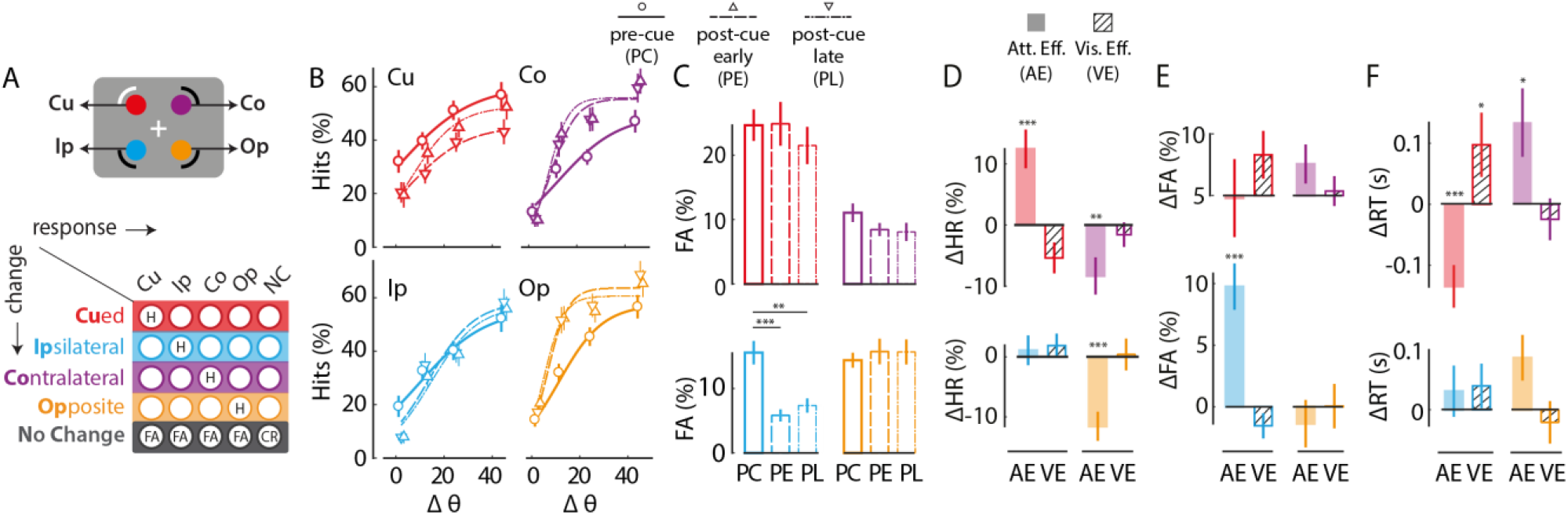
Effects of exogenous cueing on hit rates, false alarm rates, and reaction times. **A**. (Top) Schematic representation of the four locations relative to the exogenously cued location. Red: cued location (Cu); cyan: cue-ipsilateral location (Ip); purple: cue-contralateral location (Co); orange: cue-opposite location (Op). (Bottom) 5X5 stimulus-response contingency table in the 4-ADC task. Rows represent locations of change and columns represent locations of response, both measured relative to the cue. Color conventions are as in the schematic. Gray: No-change (NC) trials. H: Hit, FA: False Alarm, CR: Correct Rejection. **B**. Psychometric function (data pooled across n=42 subjects) showing hit rates for different magnitudes of grating orientation change (Δθ) for Cu (upper left panel), Ip (lower left panel), Co (upper right panel) and Op (lower right panel). Circles and solid lines: Data and sigmoid fit for pre-cue (PC) trials. Inverted triangles and dashed lines: Data and sigmoid fit for early post-cue (PE) trials. Upright triangles and dash-dot lines: Data and sigmoid fit for late post-cue (PL) trials. Color conventions are as in panel A. Error-bars: s.e.m. **C**. False-alarm rates for the four locations (Cu, Ip, Co, Op) and three trial types (PC, PE, PL). Other conventions are as in panel B. **D**. Modulation of hit-rates (averaged across orientation change values) by exogenous cueing for the four locations (Cu, Ip, Co, Op). Filled bars: Attention effect (AE=m_PC_-m_PE_). Hatched bars: Visual effect (VE=m_PE_-m_PL_). Asterisks: significance levels (*−p<0.05, **−p<0.01, ***−p<0.001). Other conventions are as in panel B. **E**. Same as in panel D, but for false-alarm rates. Other conventions are as in panel D. **F**. Same as in panel D, but for reaction times. Other conventions are as in panel D.

We quantified the attentional and visual interference effects of exogenous cueing as follows (Fig. 1B). The PC, PE and PL trial types were matched in terms of the net visual input (but not the timing) associated with the exogenous cue. We expected that pre-cueing, in the PC trials, would induce both attentional and visual interference effects. In contrast, we expected that post-cueing in the PE trials would induce visual interference effects matching those of the PC trials, but not attentional effects engaged by pre-cueing of exogenous attention. Finally, we expected that post-cueing in the PL trials would induce no pre-cueing attentional effects and much weaker visual interference effects, as compared to the PC (or PE) trials; the latter is expected because the cue appeared more than a 100 ms after the stimulus change, long enough for early sensory processing to be complete (39, 40). Therefore, subtracting each behavioral metric (e.g. sensitivity or bias) obtained from the m-ADC model (Fig. 1B) in the PE trials from its value in the PC trials yielded the attentional effects of exogenous cueing, at each location (AE(m)=m_PC_-m_PE_). On the other hand, subtracting the corresponding behavioral metrics across the PL and PE trials yielded the visual interference effects of exogenous cueing (VE(m)=m_PE_-m_PL_).

### Effects of exogenous cueing on psychometric functions of accuracy, and reaction times

First, we quantified exogenous attention’s effects on psychometric functions of hit rates (Fig. 2B), false-alarm rates (Fig. 2C), and correct rejection rates (SI Fig. S1E). Each metric revealed distinct patterns of benefits and costs across locations and cue conditions. On average, both hit rates and false-alarm rates were highest at the cued location (PC:HR: p=0.004; FA: p<0.001), and not significantly different across uncued locations. Cueing produced opposite patterns of modulations of hit rates at the cued (Cu) and two uncued (Co, Op) locations: while cueing increased hit rates significantly at the cued location in PC, as compared to PE trials (Cu: AE(hits) = 12+/−3.2%; mean+/−standard error, p<0.001 Fig. 2D), it decreased hit rates significantly at the cue-contralateral uncued (Co and Op) locations (Co_av_: AE(hits)= −10+/−2.3%, p<0.001). In contrast, cueing did not produce significant changes in false alarm rates at any of these locations (Cu: AE(false-alarm) = −0.2+/3.2%, p=0.98; Co_av_: AE(false-alarm) = 0.6+/−1.3%, p=0.53 Fig. 2E).

A notable exception to all of these trends was the cue-ipsilateral (Ip) location. In contrast to the other uncued locations, cueing did not significantly modulate hit rates at the Ip location (Ip: AE(hits) = 1+/−2.5%, p=0.38). On the other hand, it produced a striking increase in false alarm rates in PC trials, at this location (Ip: AE(false-alarm) = 10+/−1.9%, p<0.001 Fig. 2E). In addition, correct rejection rates were significantly lower in PC trials as compared to PE trials (AE(correct-rejection) = −11 +/−4.0%, p=0.01 SI Fig. S1F).

Exogenous cueing produced similar, mixed patterns of attentional effects on reaction times (RT), across cued and uncued locations. Cueing exerted effects on reaction times that closely matched the effects on hit rates. Cueing decreased reaction times significantly at the cued location in the PC, as compared to PE, trials (Cu: AE(RT_hits_) = −130+/−40 ms, p<0.001 Fig. 2F), and increased reaction times significantly at the uncued (Co_av_: AE(RT_hits_) = 110+/−30 ms, p<0.001). Again, the cue-ipsilateral location was an exception to these trends. Cueing did not significantly modulate reaction times at the Ip location (Ip: AE(RThits) = 30+/−40 ms, p=0.25 Fig. 2F).

Finally, we measured the visual interference effects of exogenous cueing, as quantified by the difference in metrics across PE and PL trials. Cueing produced a weak, suppressive visual effect on hit rates (Cu: VE(hits) = −5+/−2.5%, p=0.055), and an increase in reaction times at the cued location (VE(RT) = 100+/−52 ms p=0.018 Fig. 2D, 2F), but not at the other locations (VE(hits) p=0.95; VE(RT) p=0.88). Nor was there an effect of visual interference on correct rejection rates (p=0.48) or false alarm rates at any location (p=0.56).

Taken together, these results indicate that exogenous cueing benefitted performance in terms of increased hit rates and faster reaction times at the cued location. Corresponding costs at the uncued locations followed a striking, complementary pattern across visual hemifields: Cueing reduced hit rates and increased reaction times at locations contralateral to the cue, whereas it increased false-alarms, but did not alter hit rates or reaction times, at the location ipsilateral to the cue. We sought to synthesize these interesting, but disparate findings using the framework of signal detection theory.

### Effects of exogenous cueing on perceptual sensitivity

We asked if analyzing the data into sensitivity and bias components would resolve these mixed effects of exogenous cueing on behavioral metrics. To estimate sensitivity and bias from the 5×5 stimulus response contingency table, we employed a multidimensional signal detection model, the m-ADC model (Fig. 3A) (36, 38). The model is parameterized by sensitivity and criterion parameters, one set at each location (see Methods). Sensitivity (d’) at each location represents the discriminability of signal from noise (change versus no change) at each location. On the other hand, the criterion (c) at each location, represents a detection threshold for sensory evidence for reporting a change at each location (for a full description, see Methods).

**Figure 3.**
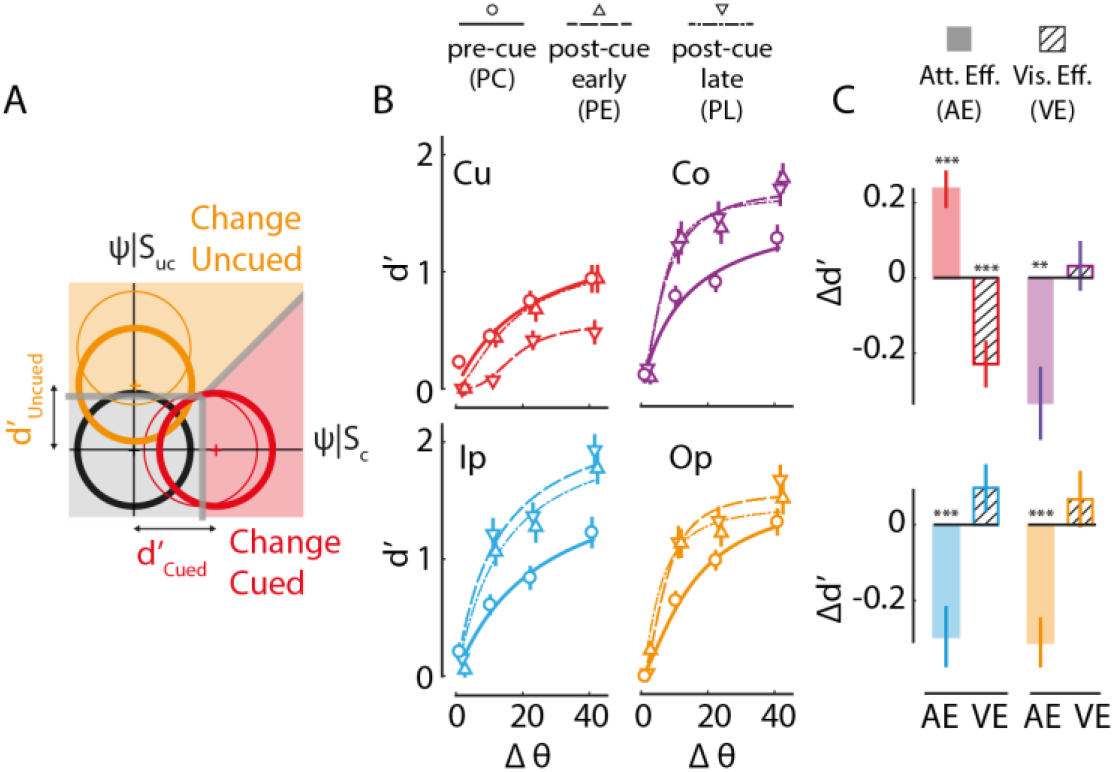
Effects of exogenous cueing on perceptual sensitivity. **A**. Schematic of m-ADC model depicting exogenous cueing effects on perceptual sensitivity (d’). Thin circular contours represent multivariate decision variable distributions whose means correspond to sensitivity values at baseline; thick contours represent changes in sensitivity due to exogenous attention at the cued and uncued locations. Other conventions are as in Fig. 1C. **B**. Psychophysical functions (data pooled across n=42 subjects) showing sensitivity, estimated from the m-ADC model, at different orientation change magnitudes (Δθ) for the four locations (Cu, Ip, Co, Op) and three trial types (PC, PE, PL). Curves: Naka-Rushton fit. Other conventions are as in Fig. 2B. **C**. Modulation of sensitivity (averaged across orientation change values) by exogenous cueing for the four locations (Cu, Ip, Co, Op). Filled bars: Attention effect. Hatched bars: Visual effect. Other conventions are as in Fig. 2D.

Before estimating cueing effects on sensitivity and bias, we tested the validity of the model for explaining subjects’ behavioral data in this four-alternative task. We performed a randomization goodness-of-fit test, based on the chi-squared statistic (see Methods). The goodness-of-fit tests yielded p-values that ranged between 0.62 and 0.99 (mean+/−stderr = 0.86+/−0.08) across subjects indicating that the model was able to fit the data successfully (SI Fig. S1H-J).

We examined the effect of exogenous cueing, first, on perceptual sensitivity (this section) and, next, on choice bias (next section). First, we quantified exogenous attention’s effect on sensitivity (AE(d’)). Exogenous cueing improved perceptual sensitivity at the cued (Cu) location (Fig. 3B-C): d’ was significantly higher in the PC trials, as compared to the PE trials, mimicking the increase in hit rates (Cu: AE(d’) = 0.24+/−0.05, p<0.001). However, in stark contrast to the mixed effects on hit rates (Fig. 2D), exogenous cueing reduced sensitivity uniformly, at all uncued locations (Ip, Co, Op): d’ was significantly lower in the PC trials (Fig. 3B, solid lines) as compared to PE trials (Fig. 3B-C; Uc: AE(d’) = −0.31+/−0.07, p<0.001). The magnitude of the attention effect of cueing was not significantly different across uncued locations (p=0.902).

Next, we quantified the visual interference effect of exogenous cueing on sensitivity (VE(d’) Fig. 3C). We found a strong reduction in d’ indicating suppression of sensory information processing by the cue: average sensitivity in the PE trials as significantly lower than that in the PL trials (Cu: VE(d’) = −0.23+/−0.06, p<0.001). In contrast, at all other uncued locations, visual interference effect of the cue was minimal or absent (Uc: VE(d’) = 0.07+/−0.04, p=0.123). In fact, the magnitude of the local visual suppression was so large at the cued location that sensitivity was, overall, lowest at the cued location as compared to all other uncued locations in the pre-cue trials (SI Fig. S2D, p<0.001).

We asked whether these effects of exogenous cueing on d’ depended on cue contrast. Specifically, we hypothesized that cues of lower contrast would produce a weaker visual suppression effect. One-third (14/42) of our participants were tested with full (100%) contrast exogenous cues whereas the remaining two-thirds (28/42) were tested with half (50%) contrast cues. As hypothesized, visual interference produced a strong and significant reduction in d’, at the cued location alone, when the cue was full contrast (Cu: VE(d’) = −0.31+/0.08, p=0.004, Uc_av_: VE(d’) = 0.08+/−0.07, p=0.27, SI Fig. S2A), but produced a much weaker reduction in d’ for half contrast cues (Cu: VE(d’) = −0.19+/−0.08, p=0.020, Uc_av_: VE(d’) = 0.06+/−0.05, p=0.29, SI Fig. S2B). Interestingly, the two types of cues induced complementary patterns of attentional benefits and costs across cued and uncued locations. Full contrast cues produced a weak improvement in d’ at the cued location (Cu: AE(d’) = 0.15+/−0.05, p=0.012, SI Fig. S2A) but a strong reduction in d’ at all uncued locations (Ip: AE(d’) = −0.5+/−0.1, p=0.001, Co: AE(d’) = −0.6+/−0.1, p<0.001, Op: AE(d’) = −0.7+/−0.1, p<0.001). On the other hand, half-contrast cues produced a strong improvement in d’ at the cued location (Cu: AE(d’) = 0.28+/−0.07, p<0.001, SI Fig. S2B), but only a weak reduction in d’ at uncued locations (Ip: AE(d’) = −0.20+/−0.09, p=0.009, Co: AE(d’) = −0.2+/−0.1, p=0.20, Op: AE(d’) = −0.12+/−0.06, p=0.06).

The global attentional effect, and the local visual suppression effect, of exogenous cueing, were supported by two other lines of evidence. First, a split-half analysis of the data (Methods) revealed that modulations of sensitivity due to exogenous attention showed a trend toward negative correlations across cued and uncued locations (AE(d’): ρ=−0.30, p=0.05), indicating a globally conserved neural resource for sensory processing (SI Fig. S2C). In contrast, no such correlation was observed for modulations of the visual interference effect (VE(d’): ρ=−0.14, p=0.39). Second, d’ values between the cued and uncued locations were significantly correlated, across subjects, in the PL trials and in PC trials (PL: ρ=0.54; p<0.001, PC: ρ=0.44; p=0.004; SI Fig. S2E), but not in the PE trials (PE: ρ=0.22, p=0.17). These results can be explained as follows: At baseline (PL trials), average perceptual sensitivities were similar across locations for each subject and, therefore, correlated across locations. Pre-cueing (PC trials) did not alter these correlations significantly, because attentional effects of exogenous cueing on d’ operated in a coupled manner (albeit with opposite signs) across cued and uncued locations. In contrast, in PE trials, the visual suppression effect of cueing on d’ were isolated to the cued location, and did not affect d’ at uncued locations, thereby eliminating the correlations across these locations.

To summarize, exogenous cueing produced two distinct effects on d’: i) a global attentional effect involving a benefit in d’ at the cued location and a corresponding cost at all uncued locations, and ii) a local, visual effect involving suppression of d’ at the cued location alone. The latter effect was mitigated somewhat with cues of lower visual contrasts. Taken together, these results indicate that exogenous attention produced a global reallocation of sensory processing resources, from uncued locations toward the cued location.

### Effects of exogenous cueing on choice bias

Next, we investigated the effect of exogenous cueing on choice bias, which we quantified with the choice criterion (b_cc_) (Methods; Fig. 4A). The value of the choice criterion has an inverse relation with choice bias: higher the choice criterion value at a location, lower is the subject’s bias for reporting a change at that location.

**Figure 4.**
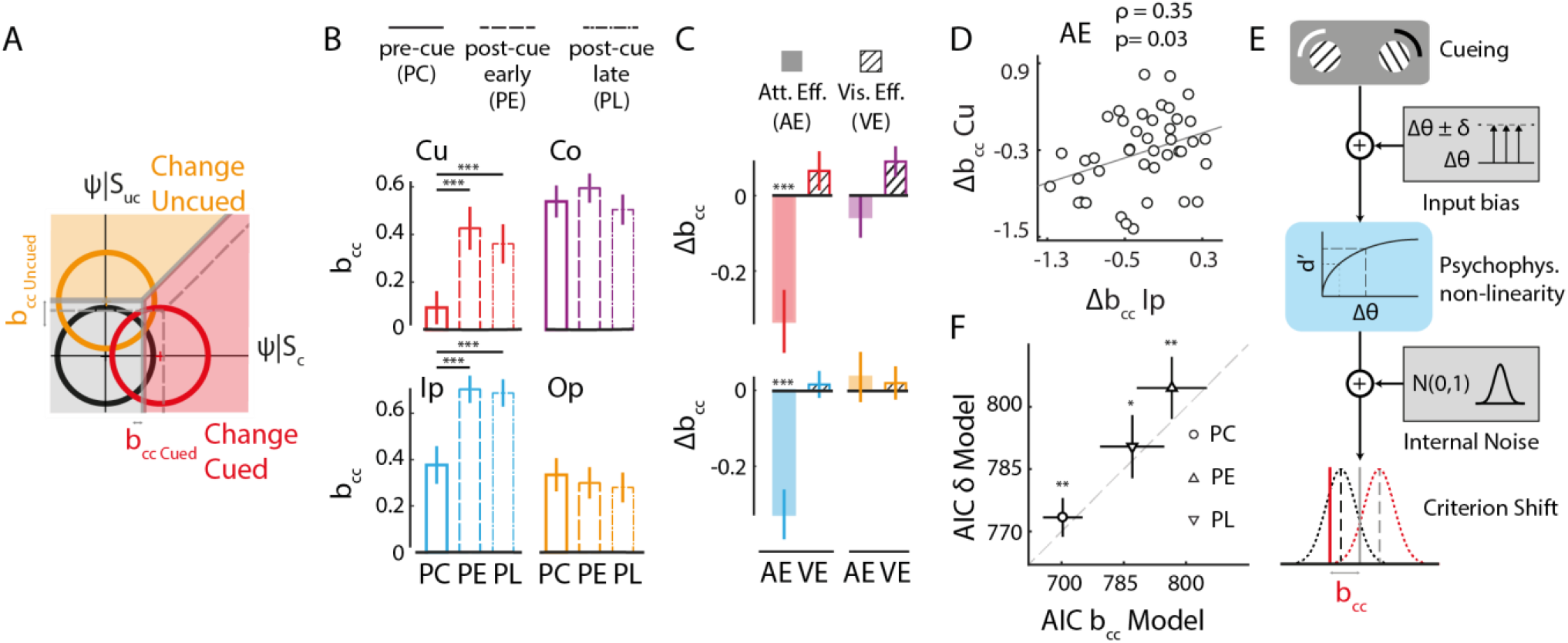
Effects of exogenous cueing on choice bias. **A**. Schematic of m-ADC model depicting exogenous cueing effects on choice bias (bcc). Dashed gray lines: Decision boundaries at baseline. Solid gray line: Change in decision boundaries due to exogenous attention. Other conventions are as in Fig. 1C. **B**. Choice bias for the four locations (Cu, Ip, Co, Op) and three trial types (PC, PE, PL). Other conventions are as in Fig. 2C. **C**. Modulation of choice bias by exogenous cueing for the four locations (Cu, Ip, Co, Op). Filled bars: Attention effect. Hatched bars: Visual effect. Other conventions are as in Fig. 2D. **D**. Covariation between attentional modulations of choice bias (AE(b_cc_)) at the cued (Cu) and cue-ipsilateral (Ip) locations. Circles: data representing individual subjects; line: best linear fit. **E**. Schematic of the input gain model, in which exogenous cueing effect on bias arises from a gain in sensory input (Δθ±δ; upper right inset) and choice bias model, in which the effect arises from a shift in the choice criterion (bcc; lowermost panel) (see text for details). **F**. Akaike Information Criterion (AIC) for the input gain model (y-axis) versus the choice bias model (x-axis). Circle, upright and inverted triangles: AIC values for PC, PE and PL trials. Error-bars: s.e.m. across subjects. Dashed gray line: line of equality.

First, we quantified the effect of exogenous attention on bias (AE(b_cc_)). Exogenous cueing significantly increased bias at the cued (Cu) location (Fig. 4B-C): b_cc_ was significantly lower in the PC trials, as compared to the PE trials (Cu: AE(b_cc_) = −0.33+/−0.08, p<0.001). On the other hand, exogenous attention produced an interesting difference in the pattern of bias changes at the uncued locations. Bias increased at the Ip location in the PC trials (Fig. 4B, solid lines) as compared to the PE trials (Fig. 4B, dashed lines), yielding a strong attention effect (Ip: AE(b_cc_) = −0.33+/−0.07, p<0.001, Fig. 4C). In contrast, no significant change in bias occurred at the other two uncued locations (Co_av_: AE(b_cc_) = −0.01+/−0.05, p=0.96). The magnitude of bias modulation was equivalent across Cu and Ip locations (p=0.94) and was significantly greater than that at either Co or Op locations (Co_av_: p<0.001). Furthermore, we found that bias modulations at the Cu and Ip locations were correlated (Fig. 4D, r= 0.35; p=0.03). These results indicate that exogenous pre-cueing induced an increase in bias, relative to post-cueing, only for locations in the hemifield ipsilateral to the cue. Moreover, the correlation in bias modulations across these locations (Cu, Ip) suggest that a common mechanism mediated the enhancement of bias across the cued-hemifield.

Second, we quantified the visual interference effect of exogenous cueing on bias (VE(b_cc_), Fig.4C). We found no measurable effect of visual interference on bias, at any location: Bias changes in PE and PL trials were not different from each other (Cu: VE(b_cc_) = 0.07+/−0.05, p=0.23; Uc_av_: VE(b_cc_) = 0.04+/−0.03, p=0.17). These results indicate that exogenous post-cueing effects on bias were not different, regardless of the latency of cue appearance, following the change.

Third, we compared the bias across locations in the pre-cueing and post-cueing conditions. We found that bias was uniformly highest at the cued location during the PC trials (p<0.001 SI Fig. S3A). In particular, the value of b_cc_ was not significantly different from zero at the cued location in pre-cueing (PC) trials (Fig. 4B). A bcc value of zero corresponds to the optimum criterion for equal prior probabilities of signal and noise, and equal rewards for each correct response. The results indicate that subjects placed their criterion close to the theoretical optimum at the cued location on PC trials (14). In contrast, in the post-cueing trials (PE, PL) b_cc_ values were significantly greater than zero at all locations. Moreover, bias was comparable across cued and opposite locations (p=0.36), but significantly lower for the ipsilateral and contralateral locations (Ip: p<0.001; Co: p<0.001). This reduction in bias across adjacent locations, as quantified by the difference between b_cc_ values at cued (Cu) and either adjacent (Ip, Co) location, were strongly correlated, both in PE and PL trials (PE: r= 0.925, p<0.001; PL: r= 0.85, p<0.001, SI Fig. S3B).

Finally, we sought to distinguish whether these choice bias modulations reflected shift in decision criteria (decisional bias) or occurred due to changes of sensory input bias or motoric response bias.

To distinguish between choice bias and sensory input bias, we fit two competing models (Methods): i) an input gain model (Model-δ), in which the criterion (c) was specified to be the same at all locations while the sensory input gain (δ) was allowed to vary across locations (Fig. 4E) and ii) a choice bias model (Model-c), in which the input gain was specified to be the same at all locations while criteria were allowed to vary across locations (Fig. 4E). We evaluated the evidence in favor of Model-δ versus Model-c based on the Akaike Information Criterion (AIC), a metric that trades off model complexity against goodness-of-fit; the model with the lowest AIC value is selected as the favored model (41). The performance of each model is shown in Figure 4F. Comparing the AIC computed by fitting each model to the data revealed that the choice bias model outperformed the input gain model (average AIC, input gain model: 789.5, choice bias model: 784.6, p<0.001, signed rank test). We conclude, therefore, that bias modulation due to exogenous cueing was a result of a shift in criterion (choice bias) rather than a change in input bias.

By task design, exogenous cues were not predictive of upcoming changes. Therefore, motoric response biases – for example, those induced by motor preparation for a response to the cued location – are unlikely to have contributed to choice bias. Nevertheless, to distinguish between choice bias and motoric response bias, we availed of the fact that greater motor bias produces faster reaction times (42). Thus, if increased motor bias contributed to increased choice bias at the cued location, we hypothesized that a) the attention effect on RT would be strongly correlated with the attention effect on b_cc_ at the cued location and b) difference in RT between cued and uncued (Co_av_) locations would also be correlated with the difference in bias (b_cc_) between these locations in the PC trials. Neither of these hypotheses were true in our data (Cu: AE(RT) vs. AE(b_cc_), ρ=0.07, p=0.65; RT: Cu-Co_av_ vs. b_cc_: Cu-Co_av_, ρ=0.16, p=0.32). These results indicate that bias modulation by exogenous cueing was not linked to motoric response biases.

We also tested whether the correlated modulation of bias (AE(b_cc_)) to the cued and cue-ipsilateral locations (Fig. 4D) occurred because of shared motor bias, because responses to changes at both of these locations (Cu, Ip) were made with fingers of the same hand (SI Fig. S1A). We hypothesized that if this were the case, we should also observe correlated modulation of reaction times at these locations (AE(RT)). Again, this hypothesis was not true in our data (AE(RT): Cu vs. Ip, ρ=0.25, p=0.12), indicating that the hemifield specific modulation of bias could not be explained by shared motor bias for responses to these locations.

To summarize, exogenous pre-cueing produced the highest choice bias at the cued location, whereas post-cueing produced a correlated decrease in bias the two locations adjacent to the cued location. The effect of exogenous attention, as quantified by the difference between pre-cueing and post-cueing effects, revealed an increase in choice bias for stimuli at locations ipsilateral (Cu, Ip), but not contralateral (Co, Op), to the exogenous cue. The increase in bias at the cued and ipsilateral locations was likely mediated by a common neural mechanism (see Discussion). Neither sensory input bias nor motoric response bias could explain these attentional effects, indicating that these reflected modulations of choice bias (shifts of choice criteria) by exogenous attention. We further tested the relationship between sensitivity and bias, and their contributions to perceptual decision dynamics, in the data.

### A synthetic model of exogenous attention’s effects on sensitivity, bias and reaction times

Exogenous attention produced a localized enhancement of perceptual sensitivity and a hemifield-wise enhancement of choice bias. These distinct patterns of sensitivity and bias enhancement indicate that exogenous cueing engaged distinct mechanisms to modulate these sensory and decisional components. To further test this hypothesis, first, we tested whether sensitivity and bias changes were correlated at each location. Sensitivity changes (AE(d’)) were uncorrelated with bias changes (AE(b_cc_)) at the cued location (ρ=−0.28, p=0.08), at the cue-ipsilateral (ρ=0.11, p=0.50), and cue-contralateral locations (ρ=-0.01, p=0.96; Fig. 5B) across subjects. Next, we tested whether modulations of sensitivity and bias were correlated across task blocks. For this, we performed a split half analysis of the data, by testing whether a change in AE(d’) was correlated with the change in AE(b_cc_), across the two halves of the data (first three versus last three blocks). No significant correlations were observed in the modulations of sensitivity and bias across blocks (SI Fig. S4A). These results indicate that distinct mechanisms mediate sensitivity and bias modulations produced by exogenous cueing (Fig. 5A).

**Figure 5.**
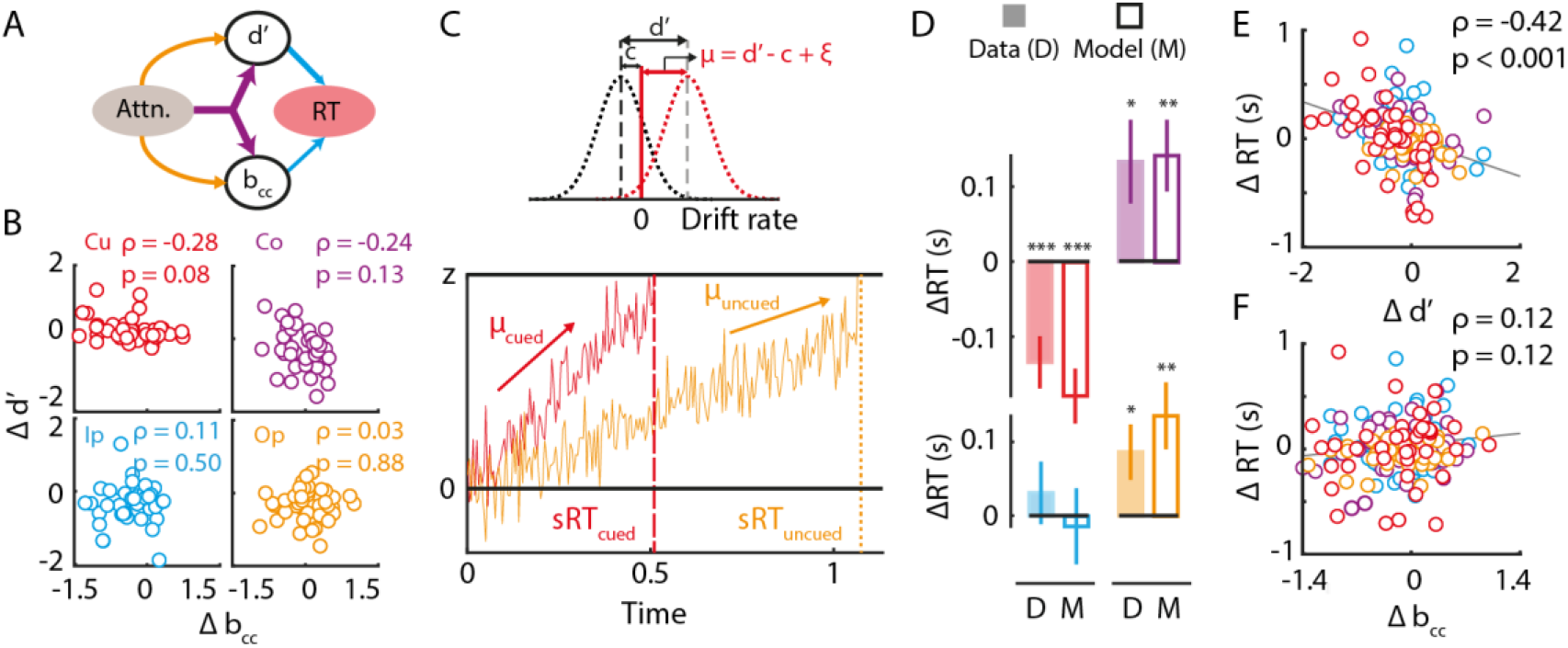
A diffusion decision model linking exogenous cueing effects on sensitivity, bias and reaction times. **A**. Schematic depicting common (purple arrows) and dissociable (orange arrows) mechanisms by which exogenous attention could modulate perceptual sensitivity (d’) and choice bias (b_cc_). Bias and sensitivity, in turn, impact reaction-times. **B**. Covariation between attentional modulations of sensitivity (AE(d’)) and bias (AE(b_cc_)) at each of the four locations. Data points: Individual subjects. ρ and p values for each correlation are indicated in the upper right corner of each sub-panel. Other conventions are as in Fig. 2B. **C**. Schematic of Diffusion Decision Model (DDM) that relates reaction times to sensitivity and bias. (Top) Schematic of relationship between drift rate (μ) in DDM and sensitivity (d’) and criterion (c) in the m-ADC model; μ = (d’ − c + ξ) where ξ is an offset parameter (see Methods). (Bottom) Simulated runs of evidence accumulation when a change occurred at the cued location (red) or at the uncued location (orange). In each case, the reaction time (sRT; dashed or dotted vertical line) corresponds to the time at which the accumulated evidence crossed a threshold or bound (z). **D**. Attention modulation of reaction times (AE(RT)) computed from model simulations (M; unfilled bars) showed close agreement with that computed from the observed data (D; filled bars), at all locations. Error-bars: s.e.m. Other conventions are as in Fig. 2D. **E**. Covariation of attentional modulation of RT (AE(RT)) with attentional modulation of sensitivity (AE(d’)). Circles: Data for each subject; colors represent values at each location. Line: best linear fit. Other conventions are as in Fig. 2A. **F**. Same as in panel E, but for covariation of attentional modulation of reaction times (AE(RT)) with attentional modulation of bias (AE(b_cc_)). Other conventions are as in panel E.

Next, we asked how these distinct sensitivity and bias modulations, each, influenced the dynamics of perceptual decisions in this task. In particular, we sought to synthetically explain, within a single framework, the disparate patterns of changes in sensitivity, bias and reaction time (RT) induced by exogenous cueing. For this, we availed of the diffusion decision model (DDM) framework, which combines signal detection theory with the drift diffusion model tying together the concepts of sensitivity, bias and reaction times in terms of a diffusion-rate and a decision threshold (43). According to DDM, decision-making is modeled as a stochastic evidence accumulation (diffusion) process. The stopping time of the evidence accumulation process corresponds to the reaction time. The drift-rate of the process captures the rate of evidence accumulation and is related to the d’ value: higher the d’ value, faster the evidence accumulation, and faster the reaction time. In our model, the criterion defines the origin of the drift rate, such that the lower the criterion, the higher the net drift rate at that location (Fig. 5C) (43, 44). Thus, the lower the decision criterion, the higher the bias, and the faster the reaction time. When the accumulated evidence reaches the decision threshold (Z), a decision in favor of reporting the signal is made. Although the DDM has not been formally extended to multialternative m-ADC tasks (44) we sought to explain qualitative trends in the data, drawing inspiration from this model.

We plotted the attention effect of exogenous cueing (AE) on sensitivity, bias and reaction times alongside (SI Fig. S4B). The data indicated that exogenous attention’s effects on in reaction-time, d’ and b_cc_ were disparate, but could be related within the DDM framework. Because cueing produced the highest gain in d’ and bias occurred at the cued location, corresponding to a higher drift rate at this location, RT was lowest at this location (SI Fig. S4B top-left). In contrast, because cueing produced a reduction in d’ (slower drift) at the Co and Op location, but produced no effect on bias, RT increased at these locations (SI Fig. S4B, right). Finally, at the Ip location, cueing produced a reduction in d’ but a gain in bias, which potentially canceled out each other’s effects, thereby resulting in no net effect on RT (SI Fig. S4B bottom-left). We tested these hypotheses by performing simulations using the DDM model (Methods). Simulated effects of exogenous cueing, based on the DDM, were strikingly similar to those observed in the data, confirming a putative mechanistic link between sensitivity, bias and RT effects of exogenous cueing (Fig. 5D).

We tested which of the parameters, d’ or bias, produced a greater effect on RT in the data. We observed that AE(RT) was significantly correlated with AE(d’) (ρ=-0.42, p<0.001) but not AE(b_cc_) (ρ=0.12, p=0.12; Fig. 5E-F). Next, we fit a multiple linear regression model with AE(RT) as the dependent variable and AE(d’) and AE(b_cc_) as the independent variables; we performed this analysis by normalizing the sensitivity and bias predictors (Methods) so that the magnitude of the standardized linear coefficients β_d_ and β_cc_ could be directly compared. We found that while β_d_ was negative and significantly different from zero (β_d_=-0.090, p<0.001), β_cc_ was positive, but not significantly different from zero (β_cc_=0.021, p=0.81; SI Fig. S4C). Similarly, an ANOVA analysis revealed a main effect of sensitivity (AE(d’): p=0.002) on reaction time (AE(RT)), but no main effect of bias (AE(b_cc_): p=0.27); the interaction between sensitivity and bias effects was also not significant (p=0.435). Finally, we performed a prediction analysis with a leave-one-out approach: we estimated β_d_ and β_cc_ on all but one subject in the population, and then predicted the reaction time effect in the left-out subject based on these population β-s and the subject’s own sensitivity and bias (Methods). This analysis revealed robust prediction accuracies (ρ=0.42, p<0.001) across the population of subjects. In addition, AE(d’) predictors were more important than AE(b_cc_) predictors for generating successful RT predictions (p=0.002, jackknife resampling; Methods).

To summarize, simulations showed that exogenous attention’s effects on reaction time could be qualitatively explained through the DDM model, linking reaction times with sensitivity and bias. Quantitatively, exogenous attention’s effects on sensitivity, rather than bias, were strongly predictive of benefits on reaction times (Fig. 5A). Taken together these results provide a synthetic model of sensitivity and bias effects on reaction times, and further confirm the dissociation between sensitivity and bias effects induced by exogenous cueing.

## Discussion

When a salient stimulus automatically captures our attention, does it enhance the perception of stimuli at the attended location, does it provide higher decisional priority for stimuli at that location, or both? To answer this question, we designed a multi-alternative change detection task with exogenous cueing, and analyzed our data with a multidimensional signal detection model. Our results reveal that exogenous attention systematically enhanced both sensory processing (perceptual sensitivity) and decisional priority (choice bias) for stimuli at the exogenously cued location.

### Dissociable effects of exogenous cueing on sensitivity and bias

Exogenous attention produced a gain in perceptual sensitivity at the cued location accompanied by an associated cost at all uncued locations, suggesting a flexible redistribution of sensory processing resources at the rapid (~100 ms) timescales of exogenous cueing. Exogenous attention produced the highest bias at the cued location (Fig. 4B), enhanced bias at both the cued location and the cue-ipsilateral location but produced no comparable change at both cue-contralateral locations (Fig. 4C). Sensitivity and bias modulations induced by exogenous cueing were uncorrelated at each location. Visual interference due to the cue produced a localized suppression of sensitivity (but not bias) at the cued location alone, in a manner that depended on the contrast of the cue. Interestingly, these robust spatial effects of cueing occurred although subjects were fully aware that the exogenous cue’s location was irrelevant to detecting the change. These results are in line with earlier studies, which have demonstrated that exogenous cueing engages attention in an automatic and involuntary manner (23, 45).

These effects of exogenous cueing were revealed by analyzing behavior in a multialternative detection task with a multidimensional signal detection model: the m-ADC model (36). Our choice of using the multialternative task and m-ADC model was motivated by several reasons. First, the task permitted measuring behavioral effects of exogenous cueing at multiple locations in the visual field. However, cueing-induced changes in behavioral metrics such as accuracies (hit rates), false-alarm rates and reaction times can arise due to a mixture of sensory and decisional factors. In our task, exogenous cueing effects on these behavioral metrics produced puzzling trends in cueing effects at uncued locations, which were not readily explained (Fig. 2D). By contrast, analysis with the m-ADC model permitted measuring these trends in terms of sensitivity and bias changes, for which the trends were neatly resolved (Fig. 3C, 4C). Second, our choice of a detection and localization task, rather than a response probe task (eg. (35)) was motivated by the recent literature, which suggests that bias in response probe tasks is linked to signal expectation (priors) rather than attention mechanisms (35). Moreover, in our task, the exogenous cue was not relevant for detecting and localizing the change, and subjects were made aware of this explicitly. Thus, our localization task permitted quantifying a spatial detection bias at each location, and its modulation by exogenous attention. Thirdly, the m-ADC model, in combination with the diffusion decision model (DDM) permitted synthetically explaining the mixed effects of exogenous cueing on reaction times: reaction times were faster at the cued location and slower at the contralateral locations, but did not change at the ipsilateral location. These disparate effects could be parsimoniously explained with DDM simulations, in which the drift rate was determined by a combination of m-ADC sensitivity and bias at each location. To our knowledge, our study provides the first demonstration of how exogenous attention modulates sensitivity and bias to produce systematic effects on behavioral accuracies and reaction times.

### Limitations and extensions

Despite revealing these interesting trends, our study leaves open a few questions to be addressed in future work.

First, exogenous cueing produced a localized increase in sensitivity but a hemifield-specific increase in bias; the latter was evidenced by correlated modulation of choice criteria, across the pre-cue and post-cue conditions, at the cued (Cu) and cue-ipsilateral (Ip) locations (Fig. 4D). Similar, hemifield-specific effects on biases have not been observed with endogenous cueing (38). Control analyses revealed that neither the benefits in bias at the cued location, nor the correlation of the bias between the Cu and Ip locations, could be attributed to sensory input gain or motoric response bias. Therefore, we propose that exogenous cueing produces not a unitary “attention field”, but a localized “sensitivity field” and a more diffuse “decisional bias field” (46). In a real-world setting, the diffuse decision bias could reflect the need for prioritized gating of information not only at the location of the stimulus that captured attention, but also for other stimuli in the neighborhood (same visual hemifield) as the attended stimulus. This hypothesis, and its neural substrates, must be investigated in future studies.

Second, a key aspect not examined by our study was the time course of sensitivity and bias enhancements induced by exogenous cueing. Previous studies have reported an inhibition-of-return (IOR) effect that occurs when the discriminandum (or change event) occurs at extended delays (>200 ms) following the cue (47, 48). It would be interesting, in future work, to test whether the localized increase in sensitivity at cued location and cue-ipsilateral increase in bias are both affected by the inhibition-of-return effect. The answers would have important implications for understanding whether the inhibition-of-return emerges from a sensory processing deficit, a change in decisional gating, or both.

### Exogenous and endogenous attention: shared or distinct neural substrates?

Currently, there is little consensus about whether exogenous and endogenous cueing of attention share the same or different neural substrates (49, 50). A potential reason for this is because numerous experiments have investigated the effects of attention with endogenous cueing (51, 52) but comparatively few studies have explored the effects of exogenous cueing of attention.

Previous studies that have compared endogenous and exogenous attention’s effects on behavior and their respective neural substrates have reported mixed results (1, 53). For example, a previous study suggested that exogenous attention primarily operates by enhancing both contrast and response gain while endogenous attention operates by increasing contrast gain only (54, 55). In contrast, a follow-up study showed, using a normalization model framework, that both endogenous and exogenous attention produce similar effects (contrast gain or response gain) depending on the relative sizes of the stimulus and attention field (24, 56, 57). Similarly, functional imaging studies have provided evidence for common (50), overlapping (25) or distinct (58) neural substrates of the two types of attention.

Our recent work (38) investigated the effects of endogenous cueing of attention, on sensitivity and bias with a four-alternative change detection task, with a stimulus configuration similar to the one used in the present study. Comparing exogenous and endogenous cueing effects on multi-alternative decision-making revealed many similarities but also certain key differences with regard to how the two types of attention operate.

At first glance, several important similarities were apparent. First, both exogenous and endogenous attention produced a gain in perceptual sensitivity, which was spatially localized to the cued location. In each case, perceptual sensitivity was modulated in a response gain-like manner, consistent with a smaller attention field compared to stimulus size. Second, the bias induced by pre-cueing was the highest at the cued location for both forms of attention. Third, we found that both exogenous and endogenous attention produced decisions that were more optimal at the cued location. In the case of endogenous cueing, indices of optimal decision-making were systematically higher at the cued location (38), while in the case of exogenous cueing, optimal values of choice criteria occurred, specifically, at the cued location. Finally, both endogenous and exogenous attention produced uncorrelated modulations of sensitivity and bias, indicating that sensitivity and bias effects are dissociable and likely to be mediated by distinct mechanisms, for both types of attentional cueing.

Nevertheless, several key differences were also observed. First, exogenous cueing of attention produced a strong cost (reduction) for perceptual sensitivity at the uncued locations relative to baseline; the same was not observed with endogenous cueing (38). Second, we also found that while exogenous cueing modulated attentional bias in a hemifield specific manner, primarily producing an increase in bias at the locations ipsilateral to the cued hemifield, endogenous cueing modulated bias in a graded manner that depended on cue validity. Third, reaction time effects of exogenous cueing were strongly correlated with sensitivity, rather than bias, whereas the reverse was true for reaction time effects of endogenous cueing (38).

In summary, our study shows that exogenous attention, does not represent a unitary phenomenon, but rather reflects the operation of distinct perceptual sensitivity and choice bias processes. The results motivate the search for the specific neural correlates of these sensitivity and bias processes. Moreover, these findings suggest that while exogenous and endogenous cueing may engage some shared mechanisms, there are also likely to be significant differences. Therefore, rather than investigating common or distinct neural substrates of exogenous and endogenous attention, it may be more fruitful to investigate whether neural substrates of each component (sensitivity and bias) are shared across the two types of attention.

## Materials and Methods

### Participants

50 subjects (28 males, age range 18-43 yrs, median age 23 yrs) with no known history of neurological disorders, and with normal or corrected-normal vision participated in the experiment. All participants provided written informed consent, and experimental procedures were approved by the Institute Human Ethics Committee at the Indian Institute of Science, Bangalore. Five subjects were excluded from analysis because of improper gaze fixation (see section on Eye Tracking). Three more subjects were excluded due to poor behavioral performance at the uncued locations (d’<0.25) in post-cued conditions. Data from 42 subjects were included in the final analysis.

### Task

Participants were tested on a cued, five-alternative change detection task. Subjects were seated in an isolated room, with head positioned on a chin-rest 60 cm from the center of a contrast calibrated visual display (22-inch DELL LCD monitor; contrast calibrated with an X-Rite i1 Spectrophotometer). Stimuli were programmed with Psychtoolbox (version 3.0.11) using MATLAB R2016b (Natick, MA). Responses were recorded with an RB-840 response box (Cedrus Inc). Fixation was monitored with a 60 Hz infrared eye-tracker (Gazepoint GP3).

We employed a previously published 4-ADC task design (37, 38), additionally, with exogenous cueing. Subjects began the task by fixating on a fixation cross (0.5° diameter) at the center of a grey screen. 1500 ms after fixation cross onset, four full-contrast, sine-wave gratings appeared, one in each visual quadrant at a distance of 5° from the fixation cross (grating diameter: 2° visual angle, spatial frequency, 3.5 cycles/degree). The orientation of each grating was drawn from a uniform random distribution, independent of the other gratings, and pseudorandomized across trials. Concurrently with the grating stimuli 4 cue placeholders (dark, full negative contrast, arcs) were presented each adjoining one stimulus. The center of each placeholder was at ±45° relative to the horizontal, and 6.7° visual angle from the fixation cross center. Each arc subtended 1.3° visual angle, with a thickness of 0.4° visual angle. The center of each placeholder was separated from the edge of the adjacent grating by a distance of 0.9° visual angle (Fig. 1A).

After a variable delay following stimulus presentation (600 ms-2200 ms, drawn from an exponential distribution), in one-third of the trials (pre-cue or PC trials), the contrast of one of the 4 placeholders (randomly chosen) was transiently increased (for 50 ms) to a high value (50% or 100% positive contrast, see below), and reverted to baseline, mimicking a bright flash; this flash corresponded to the exogenous cue. After 17 ms, the gratings were blanked for 50 ms following which they reappeared. Following reappearance, either one of the four gratings had changed in orientation, or none had changed. Subject indicated the location of change (one of four locations), or not having detected the change at any location, by pressing one of five buttons on a response box (SI Fig. S1A). In the remaining two-thirds of the trials (post-cue), the cue appeared at one of two delays, either at 17 ms (“early post-cue”, PE) or 117 ms (“late post-cue”, PL) after grating reappearance. The rationale for the cue timings for these control conditions (post-cue trials) are described in the Results section on “*Quantifying attentional and visual effects of exogenous cueing with a multialternative decision task*” (see also Fig. 1B). As in the pre-cue trials, subjects provided their response, by pressing one of five buttons.

We term trials in which a change in orientation occurred at one of the four locations as “change” trials (80% of all trials), and trials in which no change in orientation occurred as “no change” trials (20% of all trials). We term the location closest to which the cue occurred as the “Cued” (Cu) location, the uncued location diagonally opposite to the cued location, as the “Opposite” (Op) location, and the two other uncued locations as “Ipsilateral” (Ip) or “Contralateral” (Co) locations, depending on whether they were in the visual hemifield ipsilateral or contralateral, respectively, to the cued location (Fig. 2A). Changes were equally and pseudo-randomly distributed among these four locations. Thus, the cue was uninformative spatially and had a conditional validity of 25% on change trials among the four locations. The experiment was run in six blocks of 60 trials each (total, 360 trials per subject), with no feedback. In a training session prior to the experiment, subjects were familiarized with the task and response protocol for 1-2 blocks (60 trials each), with explicit feedback provided at the end of each trial about the location of the change and the correctness of their response. Data from these “training” blocks were not used for further analyses.

We tested three versions of the above task protocol, with slight variations. In the first protocol, the exogenous cue was presented at full (100%) contrast with no time limit given to the subjects for response (n=14 subjects). The second variant was identical with the first protocol, except that the exogenous cue was presented at 50% contrast (n=14). The third variant was identical with the second protocol, except that subjects were required to respond within a maximum window of 1250 ms (n=14). Results were closely similar for all protocols and, unless otherwise indicated, data were pooled across protocols.

### Eye-tracking

Subjects’ gaze was binocularly tracked and the deviation in their gaze from the fixation cross was recorded and stored for offline analysis. Trials in which the eye-position deviated by more than 2 degrees radially from the fixation cross from the onset of the cue (pre-cue trials) or onset of the blank (post-cue trials) until the final response were removed from further analysis. Our subjects were of South-Asian origin and exhibited dark pigmentation of the iris, rendering it nearly indistinguishable from the pupil. Hence, the contrast of the pupil (relative to the iris) was weak, and the tracker occasionally lost the location of the pupil; trials in which this occurred for more than 100 ms continuously were also excluded from the analysis. We excluded data from subjects for whom the combined rejection rate (eye deviation and lost tracking) exceeded more than 25% of all trials (5/50 subjects). The median rejection rate for subjects included in the analysis was 8.8% [0.0-22%] (median, 95% confidence intervals). We also tested with a randomization test, based on the chi-squared statistic (see below), whether these rejected trials significantly altered the distribution of responses in the contingency table for any subject, and found that this was not the case (p-values for distributions before and after rejection: 0.99, mean across subjects).

### m-ADC psychophysical model

Previous studies have employed one-dimensional signal detection models to distinguish sensitivity from bias in detection and localization tasks (12, 59). However, such an approach represents a model misspecification for analysis of behavior in multialternative localization tasks, as elaborated in (36). We applied a recently developed multidimensional signal detection model, the m-ADC model to analyze subjects’ behavior in this 5-alternative task. The m-ADC overcomes the pitfalls of previous approaches by modeling the decision on each trial in a multidimensional decision space. Specifically, for the 4-ADC model used in this study, sensory evidence for the change event is represented along four orthogonal decision variable axes, one corresponding to each location. Decision surfaces are represented as manifolds comprising of 10 intersecting hyperplanes in a 4-dimensional space (36). The hyperplanes belong to the family of optimal decision surfaces for distinguishing signals of each class (a change event at each location) from noise (no change), and for distinguishing signals of each class from the other (a change event at one location from a change event at another location). These hyperplanes are parametrized by four criteria, which reflect detection thresholds for reporting change versus no change at each location. In addition, the model is parameterized by four sensitivity (d’) values that reflect the signal strength at each location, and are quantified as the distance between the means of the signal and noise distributions, measured in units of noise standard deviation, along the corresponding decision variable axis (Fig. 1C).

The model decouples and separately quantifies perceptual sensitivity and choice criteria at each location based on the 5×5 contingency table of responses obtained from the multi-alternative attention task (Fig. 2A, SI Fig. S1D). A description and mathematical derivation of the model are provided in (36, 37). An extension of the m-ADC model to tasks that seek to measure the entire psychophysical function at the cued and uncued locations is provided in (38), and this latter model is used for behavioral analysis in this study.

### Contingency tables

Subjects’ responses in the task were used to construct 5×5 stimulus response contingency tables, one for each of the pre- and post-cue trial types (PC, PE, PL). Change locations were represented on the rows and response locations on the columns and no-change events and responses were represented in the last row and last column respectively (Fig. 2A). The contingency table was then restructured so that all change events and responses were measured relative to the cued location. Each contingency table comprised five categories of responses: hits, misses, false-alarms, mislocalizations (or misidentifications) and correct rejections (SI Fig. S1D). Since four values of orientation changes were tested at each location, each contingency table contained 68 independent observations: 16 hits, 48 misidentifications and 4 false-alarms; the last category of responses do not depend (by definition) on orientation change magnitude.

### Psychometric functions

To compute the psychometric function (percent correct as a function of orientation change angle), we calculated the proportion of trials in which the subject detected and localized the change accurately; this was computed separately for each location (relative to the cue) and each trial type (PC, PE, PL). Percent correct values, across all angles of orientation change, were fitted with a three parameter sigmoid function to generate the psychometric function (Fig. 2B). Psychometric functions for all subjects were estimated with a set of four change angles spanning 2° to 45° ([2,12,25,45°]), presented in an interleaved, pseudo randomized order across locations. The middle two angles (12° and 25°) were tested twice as often as the extreme two angles (2° and 45°). The average psychometric curve was generated by pooling responses across all subjects and computing the above metrics for the pooled data. False alarm and correct rejection rates were calculated based, respectively, on subjects incorrect and correct responses during no-change trials. Standard error of mean was calculated and reported as error.

### Model parameter estimation and goodness-of-fit

To compute the psychophysical function (sensitivity as a function of orientation change angle), individual subjects’ response contingencies were fitted with the m-ADC model (1). Sensitivity is expected to change depending on the change angle magnitude (Δθ); hence, different sensitivity values were estimated for each change angle tested. On the other hand, the criterion at each location was estimated as a single, uniform value across change angles, as the subject could not anticipate the change angle presented on each trial. Thus, the model estimated 20 parameters (d’ values for each of the 4 locations and 4 angles, and criteria for each of the 4 locations) from 68 independent observations in the contingency table for each trial type. Sensitivities and criteria were estimated with maximum likelihood estimation (MLE), using a procedure described previously (36).

Goodness-of-fit of the model to the data was assessed using a randomization test based on the chi-squared statistic; the procedure is described in detail elsewhere (38). A small p-value (p<0.05) for the goodness-of-fit statistic indicates that the observations deviated significantly from the model fit. The median p-values for PC trials was 0.90 (range: 0.67-1.00), for PE trials was 0.89 (range: 0.69-0.99) and PL trials was 0.83 (range: 0.62-0.98) (Fig. S1B, S1C), indicating that the model fit the observations well.

### Psychophysical functions of sensitivity and bias

The psychophysical function was generated by fitting sensitivity values across angles at each location with a three-parameter Naka-Rushton function: asymptotic sensitivity (dmax) and orientation change value at half-max (Δθ50) were fit as free parameters, whereas the slope parameter (n) was fixed at 2.0 based on pilot fits to the data. As before, a single combined psychophysical curve was generated by pooling contingencies across all subjects and fitting the m-ADC model to the pooled data (Fig. 3B).

Next, we fit the m-ADC model to the contingency table from each subject to estimate sensitivity and criteria at each location and for each trial type. We then computed the mean sensitivity across change angles (d’_av_), as well as two measures of bias: the choice criterion (b_cc_ = c-d’/2), and the likelihood ratio (b_lr_: exp(d’(c − d’/2))). Both b_cc_ and b_lr_ are candidate measures for quantifying bias (38) and both exhibited similar patterns of variation for most of the analyses reported here. As a result, we report trends based only on b_cc_ in the main text; b_lr_ values are reported in the Supporting Information Fig. S3C-F.

These analyses sought to investigate the influence of exogenous cueing on psychometric (e.g. hit rates, false-alarm rates) and psychophysical (e.g. d’, b_cc_) parameters, and to dissociate the effects of attention and visual interference due to the cue. Therefore, we analyzed our data on two levels. First, we looked at how each psychophysical measure (d’ or b_cc_) varied in each condition across the four locations (e.g. Fig. 3B). We then computed the attentional modulation effects at each location by subtracting the psychophysical measure in PE trials from the corresponding measure in PC trials, and the visual interference effects by subtracting values in PE trials from the corresponding values in PL trials (see Results, section on “*Quantifying attentional and visual effects of exogenous cueing with a multialternative decision task*”). The attention effect (AE) and visual interference effect (VE) on each metric (m), are reported in the following format: AE(m)=m_PC_-m_PE_ or VE(m)=m_PE_-m_PL_; where m_PX_ represents the value of metric m in trial type PX.

To test whether the sensitivity modulation by exogenous attention at the cued and uncued locations reflected the redistribution of limited sensory processing resources, we performed a split block analysis. Subjects performed 6 blocks of 60 trials each through the experiment. We split the experiment as the first three blocks and last three blocks and used the mADC model to compute d’ and b_cc_ as described previously. d’_av_ was computed by excluding the lowest angle. Values for the first half and the second half of the data were subtracted from each other. ΔCu and ΔUc_av_ were computed for both halves and were correlated with robust (“percentage-bend”) correlations. Corresponding correlation and p-values are shown in Figure 3D. To test whether sensitivity and bias modulations were correlated, we performed a similar split half analysis except that in this case AE(d’) and AE(b_cc_) were computed for each split half of the data, subtracted across the two halves and correlated with each other (SI Fig. S4A).

### Statistical tests

Tests of significant differences in d’ (Fig. 3B) across trial types were performed using an ANOVA treating locations and trial types as discrete factors, and change angle as a continuous factor, with Tukey-Kramer correction for multiple comparisons. A similar ANOVA was performed for b_cc_ (Fig. 4B) without angles as a factor. Significance tests comparing attention or visual interference effects on different parameters for each location were performed using a Wilcoxon signed rank test with a Benjamini-Hochberg correction for multiple comparisons. Correlations across different parameter values were computed using robust correlations (“percentage-bend” correlations) to prevent outlier data points dominating the correlations (60); correlation coefficient (ρ) and significance (p) values are indicated on the corresponding figures. Unless otherwise stated, significant p-values at the successively higher levels of significance (p<0.05, p<0.01 and p<0.001) are shown with successively higher number of asterisks (* (single-star), ** (two-stars) and *** (three stars)), above the respective bar plots.

### Model comparison analysis

To determine whether the bias quantified by the m-ADC model was a choice bias (shift in choice criteria), or could be explained by sensory input gain alone we compared two models. In the first model (input gain model) input gain (δ; Fig. 4E) induced by exogenous cueing varied across locations (Cu, Ip, Co, Op) but the criterion was held uniform across locations. The input gain is modeled as an additive factor to the input (change in Δθ), which increases the gain of the input along the x-axis of the psychophysical function (Fig. 4E) (35). In the second model (choice bias model), the input gain was uniform across locations but criteria varied across locations, as in the conventional m-ADC model. In each case, models were compared with the Akaike Information Criterion (AIC) that represents a tradeoff between model complexity (the number of fitted model parameters) and goodness-of-fit (based on the log-likelihood function); a lower AIC score represents a better candidate model. The number of fitted parameters in both of the input gain and choice bias models were 10. We did this using a parameterized form of the m-ADC model using dmax (4), θ50 (1), criteria (1), δ (4) as parameters for fitting the data in Model-δ and dmax (4), θ50 (1), criteria (4), δ (1) in Model-c. AIC values were computed for each trial type and subject separately. Average AIC values with s.e.m error bars are reported. (Fig. 4F)

### Analysis of reaction times and simulations with the Diffusion Decision Model

Reaction times were quantified as the time from the onset of the change until the subject’s button press response. As before, reaction times were compared across cued and uncued locations with ANOVA and Tukey-Cramer post-hoc tests.

We simulated the effect of sensitivity (d’) and bias (b_cc_) on reaction-times (RT) using the Diffusion Decision Model (DDM; (44) which models evidence accumulation as a noisy diffusion process. In the conventional drift-diffusion model, decision time is parameterized by an accumulation rate (or drift rate, μ), starting point (origin) and stopping threshold of the evidence accumulation process (z). DDM relates parameters in the drift-diffusion model, with the sensitivity and criterion parameters of signal detection theory. Here, we used a version of the DDM in which the drift rate was modeled as a linear combination of sensitivity and criteria (44). Specifically, the sensitivity was directly related to the drift rate, and the criterion defined the origin of the drift rate at each location (Fig. 5C), such that a lower criterion corresponded to a higher, overall drift rate (μ=d’-c). Our model contained 2 additional free parameters for fitting the data: an offset term for the evidence accumulation rate (ξ=2.0) and a coefficient (k=10^−3^) for the noise term, whose values were chosen to provide close fits to the data, although the results were robust to the choice of these parameters.

The overall model can be summarized as follows:

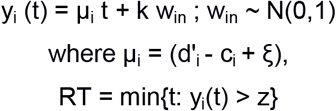

where y_i_ (t) denotes the accumulated evidence at the i^th^ location up to time t (i=1,2,3,4), μ_i_ denotes the evidence accumulation rate and win denotes the noise added at each time step (drawn from a standard normal distribution). The origin of evidence accumulation was set to zero and decision threshold (z) was set to 1 at all locations. Thus the model is akin to a “race” model with four competing evidence accumulators, one at each location, and the decision is made when one of the four accumulators crosses the decision threshold. The time step for the simulation was taken to be 1 millisecond, and accumulated evidence was calculated with the Euler method. A total of 288 trials (24 trials per location and condition) were used for the simulation, which matched our actual experimental task (only change trials were simulated). The average reaction time across trials was computed for each subject, location (Cu, Ip, Op, Co), and each trial type (PC, PE, PL), separately, based on the d’ and c for that subject, location and trial type. The attention effects and visual interference effects for the simulated RTs were then computed, with an identical procedure as for the experimental RTs (described in the Results section on “*Quantifying attentional and visual effects of exogenous cueing with a multialternative decision task*”).

### Reaction time regression and prediction analysis

To quantify the effect of sensitivity and bias on reaction times obtained from the behavioral data, we performed a multiple linear regression analysis with AE(RT) as the dependent variable and normalized AE(d’) and AE(b_cc_) as predictors. Each variable was normalized based on z-scoring to make the magnitudes of the regression coefficients, β_d’_ and β_cc_, comparable. To test if the β_d’_ and β_cc_ values were significantly different from zero we employed a random permutation test. We created two null distributions for β_d’_ and β_cc_ (each) by randomly shuffling subject labels on the predictor variables and recomputing the coefficients over 1000 iterations. The p-values correspond to the proportion of instances in the null distribution which fell below the actual β_d’_ and β_cc_ values. We also performed an ANOVA to test for the relative contributions of AE(d’) and AE(b_cc_), in explaining the variance in AE(RT).

In addition, we performed a prediction analysis, by predicting the attentional effect on reaction time for each individual subject using their sensitivity and bias modulations. For this analysis, as before, we fit AE(RT) based on AE(d’) and AE(b_cc_) as predictors using multiple linear regression. The only difference was that this fitting was done by estimating β_d’_ and β_cc_ by leaving out one subject at a time. Following this the AE(RT) of the left out subject was linearly estimated based on the AE(d’) and AE(b_cc_) of the left out subject, but using the β_d’_ and β_cc_ from the rest of the (n=41) subjects. We correlated the predicted AE(RT) against the observed AE(RT) with robust correlations (“percentage-bend” correlations). To determine which coefficient -- β_d’_ or β_cc_ -- was more important for RT predictions, we computed confidence intervals for these coefficients based on jack-knife (leave-one-out) standard error estimates (61).

## Acknowledgments

The authors would like to thank Sanjna Banerjee, Sricharan Sunder and Varsha Sreenivasan for comments on a preliminary version of this manuscript, Ambuj Misra for help data acquisition and Suhas Ganesh for help with data analysis. This research was funded by a Wellcome Trust-Department of Biotechnology India Alliance Intermediate fellowship, a Science and Engineering Research Board Early Career award, a Pratiksha Trust Young Investigator award, a Department of Biotechnology-Indian Institute of Science Partnership Program grant, and a Tata Trusts grant (all to DS).

## Supporting Information

**Figure S1.**
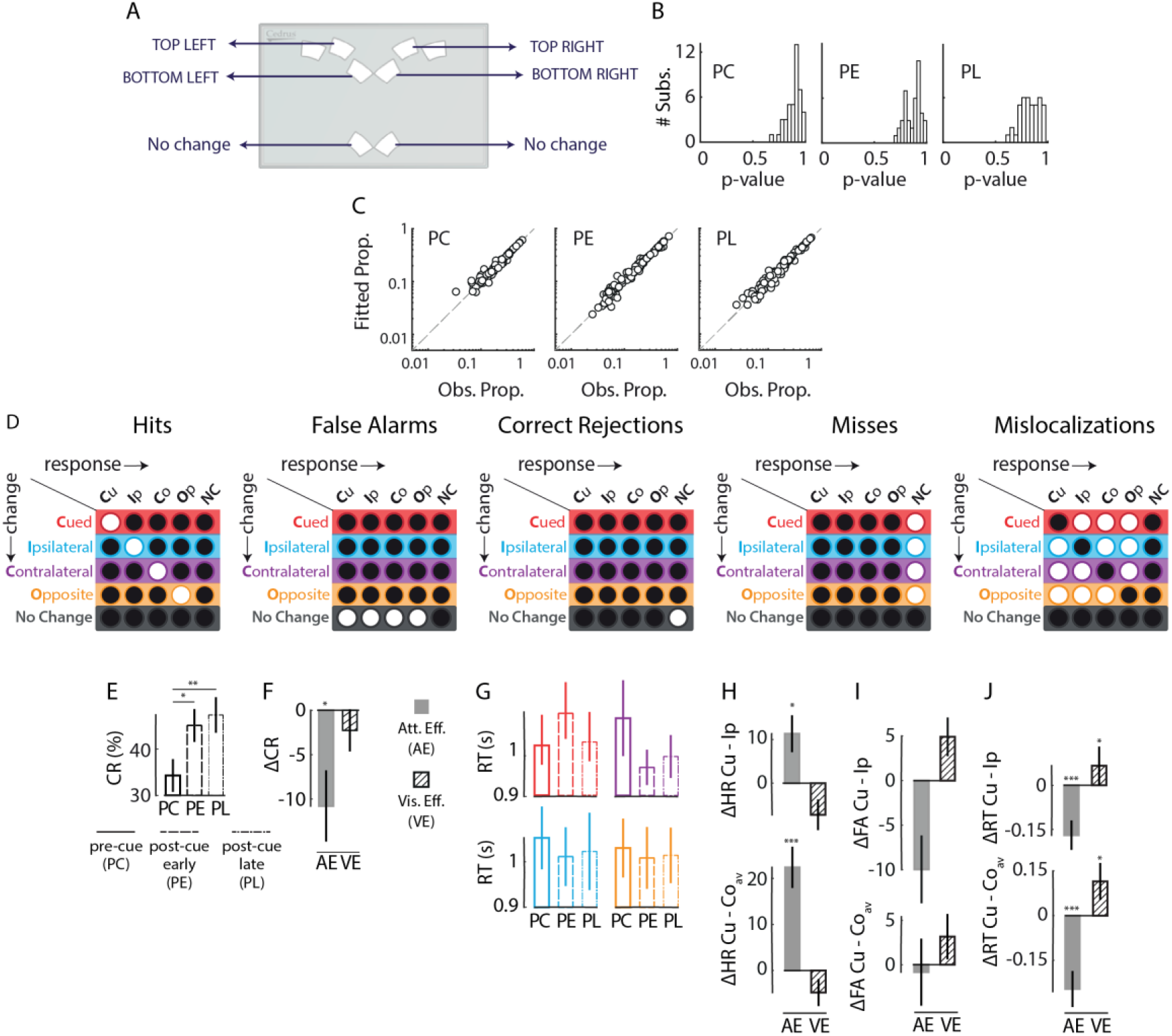
Model fitting and modulation of responses by exogenous cueing. **A**. Response box (RB-840, Cedrus Inc.) showing the configuration for responses for the 4-ADC task shown Fig. 1A (main text). **B**. Distribution of p-values for randomization goodness-of-fit tests for PC (left), PE (middle) and PL (right) trials. **C**. Response proportions fitted with the m-ADC model (ordinate) plotted against observed response proportions (abscissa) for PC (left), PE (middle) and PL (right) trials. Data points: individual stimulus-response contingencies for each of the n=42 subjects. **D**. Different response categories (highlighted circles) in the 5×5 stimulus-response contingency table. From left to right: ‘Hits’ – response indicating the correct location of change; ‘False Alarms’: response indicating change on no-change trials; ‘Correct rejections’: no-change response on no-change trials; ‘Misses’: no-change response on change trials; and ‘Mislocalizations’: response indicating change at location other than the change location. **E**. Correct rejection rates for the three trial types (PC, PE, PL). Other conventions are as in Fig. 2C (main text). **F**. Modulation of correct rejection rates with exogenous cueing. Filled bars: Attention effect. Hatched bars: Visual effect. Other conventions are as in Fig. 2D (main text). **G**. Reaction times of ‘Hits’ (averaged across orientation change values) for the four locations (Cu, Ip, Co, Op) and three trial types (PC, PE, PL). Other conventions are as in Fig. 2C (main text). **H**. (Top) Modulation of hit-rates by exogenous cueing at cued relative to cue-ipsilateral locations (Cu vs Ip). (Bottom) Same as top panel, but for cued relative to average of cue-contralateral and cue-opposite locations (Cu vs Coav). Filled bars: Attention effect. Hatched bars: Visual effect. Error-bars: s.e.m. across subjects. **I**. Same as in panel H, but for false-alarm rates. Other conventions are as panel H. **J**. Same as in panel H, but for reaction times. Other conventions are as panel H.

**Figure S2.**
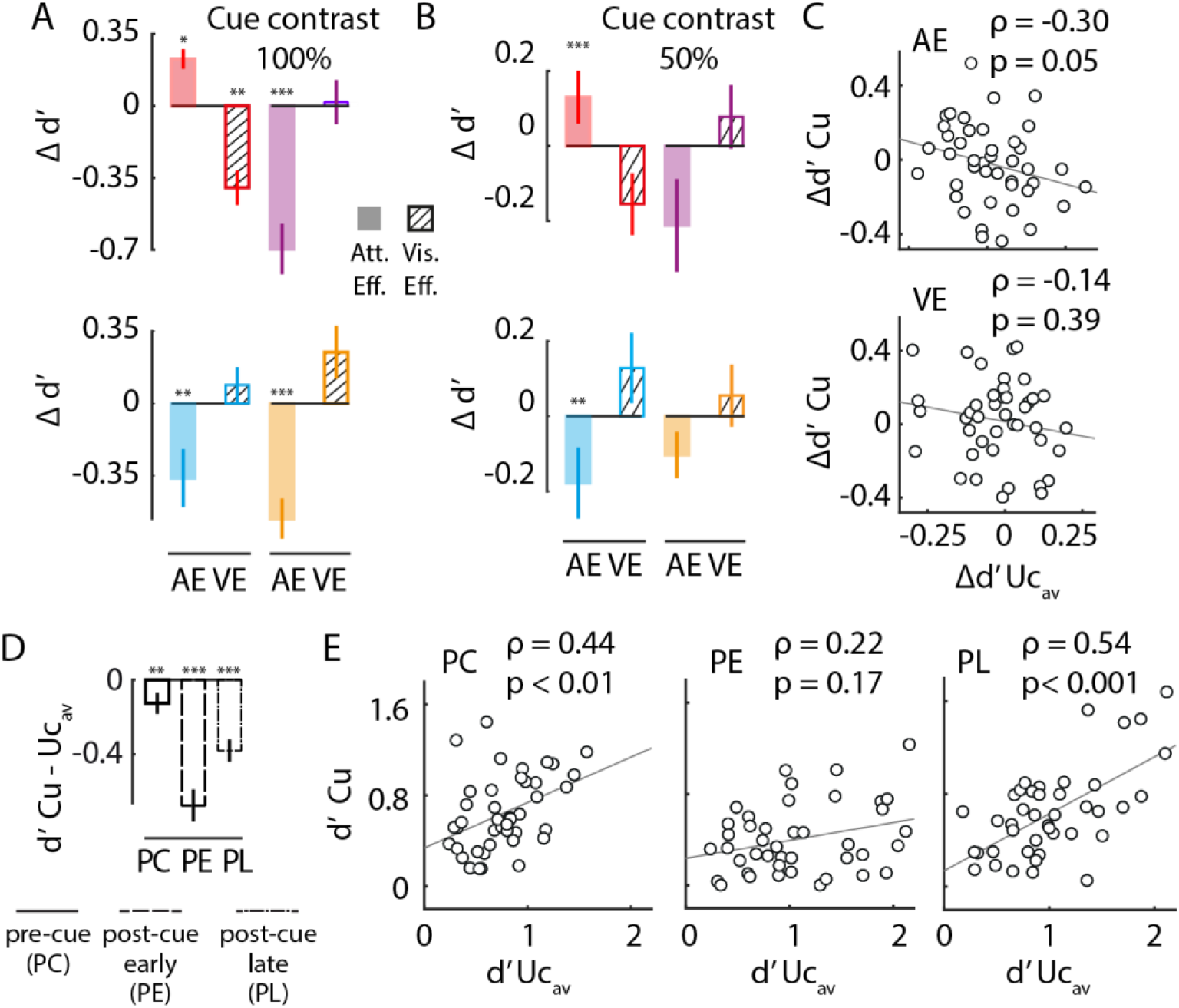
Effects of exogenous cueing on sensitivity and its modulations. **A**. Modulation of sensitivity (averaged across orientation change values) by exogenous cueing for the four locations (Cu, Ip, Co, Op) for full (100%) contrast cues (n=14 subjects). Filled bars: Attention effect. Hatched bars: Visual effect. Other conventions are as in Fig. 3C (main text). **B**. Same as in panel A but for half (50%) contrast cues (n=28 subjects). Other conventions are as in panel A. **C**. (Top) Covariation of attentional modulation of sensitivity (AE(d’)) by exogenous cueing at cued location versus at uncued locations based on split half analysis of the data (see text for details). Circles: Data for each subject (n=42). Line: best linear fit. ρ and p values for each correlation are indicated in the upper right corner of ea ch sub-panel. (Bottom) Same as top panel, but for visual effect of exogenous cueing (VE(d’)). **D**. Difference of d’ values at the cued location relative to uncued locations (average across Ip, Co, Op) for the three trial types (PC, PE, PL). Other conventions are as in Fig. S1E. **E**. Covariation of sensitivity (d’) values at cued location with uncued locations for PC (left), PE (middle), and PL (right) trials. ρ and p values for each correlation are indicated in the upper right corner of each sub-panel. Other conventions are as in panel C.

**Figure S3.**
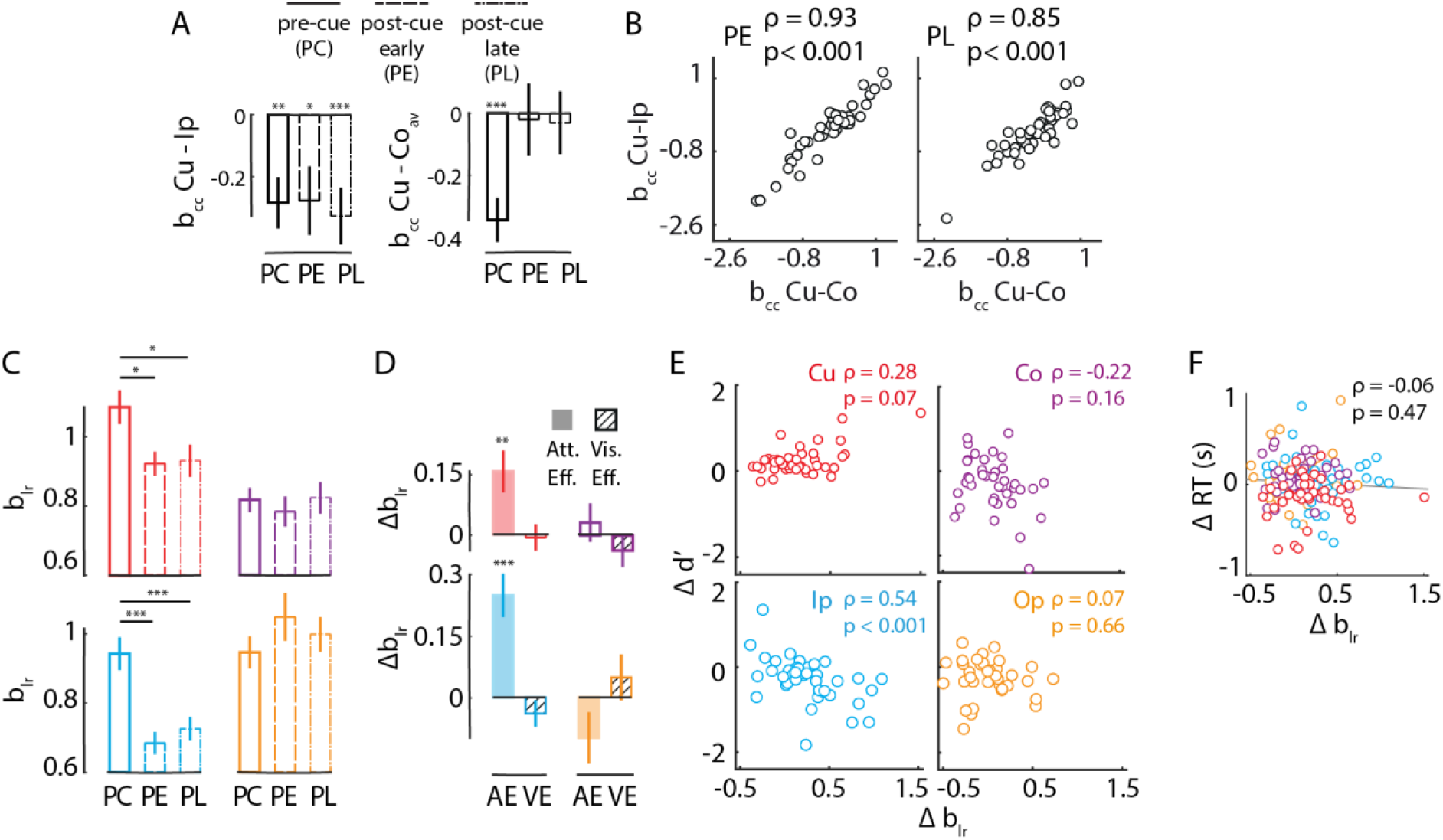
Effects of exogenous cueing on bias and its modulations. **A**. Difference of bias (bcc) values at the cued location relative to Ip location (left) and average of contralateral locations (Co, Op) (right) for the three trial types (PC, PE, PL). Other conventions are as in Fig. S2D. **B**. Covariation of difference of bias (bcc) across the cued and cue-ipsilateral location (Cu-Ip) versus the difference of values across cued and both contralateral location (Cu-Co), for PE (left) and PL (right) trials. Circles: Data for individual subjects (n=42). ρ and p values for each correlation are indicated in the upper right corner of each sub-panel. **C**. Same as Fig. 4B (main text) but for likelihood ratio bias (blr). **D**. Same as Fig. 4C (main text), but for likelihood ratio bias (blr). **E**. Covariation of modulation of sensitivity (AE(d’)) with modulation of likelihood ratio bias (AE(b_lr_)). Other conventions are as in Fig. 5B (main text). **F**. Covariation of modulation of reaction times (AE(RT)) with modulation of likelihood ratio bias (AE(br)) by exogenous cueing. Other conventions are as in Fig. 5F (main text).

**Figure S4.**
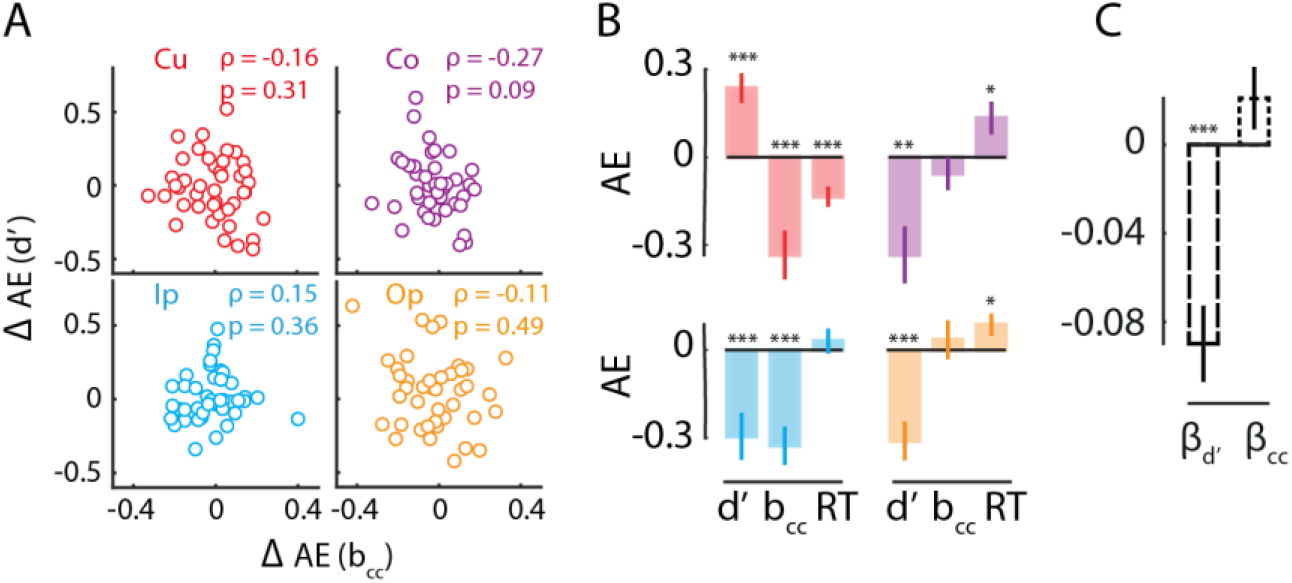
Effect of exogenous cueing on sensitivity, bias and reaction times and their modulations. **A**. Covariation between attentional modulations of sensitivity (Δ AE(d’)) and bias (Δ AE(bcc)) at each of the four locations; Δ AE(d’) and Δ AE(bcc) were computed by computing AE(d’) or AE(bcc) for each half of the data, and subtracting the corresponding quantities across the two halves. Data points: Individual subjects (n=42). ρ and p values for each correlation are indicated in the upper right corner of each sub-panel. Other conventions are as in Fig. 5B (main text). **B**. Attention effect of exogenous cueing (AE) (filled bars) on sensitivity (d’), bias (bcc) and reaction times (RT). Other conventions are as in Fig. 2D (main text). **C**. Regression coefficients for AE(d’) (β_d_) and AE(b_cc_) (β_cc_) for predicting the attentional modulation of reaction time (AE(RT)). Error bars indicate jack-knife s.e.m. Asterisks: significance levels (***−p<0.001).

